# A Systematic Benchmark for Peptide Property Prediction

**DOI:** 10.64898/2026.02.09.704773

**Authors:** Dong Xiaoying, Yang Kaijun, Wu Tianxiang, Li Pengyong, Gao Lin

**Affiliations:** School of Computer Science and Technology, Xidian University, Shaanxi, China

## Abstract

Accurate prediction of peptide physicochemical properties and biological activities is critical for rational peptide design and high-throughput screening. However, current research is often constrained by heterogeneous data sources and inconsistent evaluation standards, which hinder fair comparisons and reliable assessments of model generalization. In this work, we present PPB, a peptide property prediction benchmark designed to evaluate model performance with an emphasis on realistic generalization across both classification and regression tasks. By applying unified biological filtering criteria, we systematically curated and standardized 15 datasets comprising 161,571 unique sequences, spanning a wide range of physicochemical properties and functional activities. We benchmarked seven representative architectures—encompassing traditional machine learning, deep learning, and pre-trained language models—alongside diverse feature encoding schemes. Furthermore, we investigated the impact of random versus homology-based (sequence similarity) data splitting strategies on model robustness. To facilitate community access, we developed the PPB web server (http://ppb.molmatrix.com/index.html), which provides centralized resources for standardized dataset downloads, interactive visualization of benchmark results, and detailed evaluation protocols.

**Author summary:** Peptides are short amino acid chains essential for biological functions and drug discovery. While AI models have accelerated peptide property prediction, the field lacks a unified standard to fairly compare these methods, often leading to inconsistent results and overoptimistic performance estimates. In this study, we introduce the Peptide Property Benchmark (PPB), a comprehensive framework featuring 15 standardized datasets and over 160,000 sequences. We systematically evaluated diverse AI paradigms, including traditional machine learning and advanced protein language models. Our results demonstrate that large-scale pre-trained models—the biological equivalent of large language models—offer superior accuracy and stability, particularly for small or complex datasets. Crucially, our analysis reveals a “clustering bottleneck”: standard tools used to group proteins based on similarity often fail when applied to short peptides, causing data to fragment excessively. This suggests that traditional strategies for testing model generalization may be less effective for peptides than previously assumed. To facilitate community progress, we provide an online platform for standardized data and evaluation. This work establishes a rigorous foundation for developing more reliable AI tools for the next generation of peptide-based therapeutics.

## Introduction

Peptides typically consist of 2–50 amino acid residues and occupy an intermediate position between small molecules and proteins, playing essential roles in a wide range of biological processes [1, 2], including signal transduction, immune regulation, antimicrobial defense, and cellular transport [3, 4]. Owing to their high biological specificity, favorable structural designability, and relatively low synthesis cost, peptides have become important research targets in drug discovery, biomaterials engineering, and therapeutic molecular design. In practical applications, key characteristics of peptides—such as biological activity, toxicity, hemolytic activity, and various physicochemical properties—directly determine their functional performance and clinical translation potential.

Due to the high cost and low throughput of experimental measurements, computational approaches for peptide property prediction have gradually emerged as important tools to support peptide screening and rational design [5–14]. In recent years, a variety of predictive models have been proposed, including traditional machine learning methods based on handcrafted features [15], end-to-end deep learning models [16], and, more recently, pretrained protein language models. Despite continuous methodological advances, substantial differences remain across studies in terms of data construction, experimental settings, and evaluation protocols, making direct and reliable comparison of model performance difficult.

Existing studies often focus on a single prediction task or a specific dataset and adopt individually defined data construction and validation procedures. This practice limits fair comparisons among different models and weakens the overall understanding of their true generalization ability. At the data level, peptide property prediction tasks face significant challenges arising from data heterogeneity [17]. Publicly available peptide datasets are typically integrated from different databases and literature sources, lacking unified standards in data collection procedures, biological screening criteria, and experimental backgrounds. Moreover, differences in negative sample construction strategies across datasets for different peptide properties further introduce data noise and systematic bias, thereby significantly affecting the reliability of model performance evaluation and their assessed generalization capability.

The choice of evaluation strategy is also a critical factor influencing the assessment of model generalization. In sequence modeling tasks, a large number of highly similar sequences often exist between the training and testing sets, causing random splitting strategies to overestimate predictive performance [18]. To mitigate this issue, sequence similarity–based splitting strategies are commonly adopted, in which the similarity between training and testing sets is explicitly constrained, thereby enabling a more realistic evaluation of a model’s true predictive ability on unseen peptides. However, despite being more consistent with practical application scenarios in theory, similarity-based splitting strategies still face several underexplored challenges in the field of peptide property prediction. For example, there is a lack of consensus on similarity thresholds and clustering parameters across studies, and the sensitivity of the resulting data partitions to hyperparameter choices has not been systematically evaluated. In addition, whether the resulting clusters are reasonable in terms of size balance and representativeness remains insufficiently quantified and compared. Consequently, the effectiveness and applicability of similarity-based splitting strategies for peptide property prediction tasks require further systematic investigation.

Based on these considerations, we present the Peptide Property Prediction Benchmark (PPB), a systematic evaluation framework designed to establish a rigorous and unified standard for the field. Our work provides a comprehensive assessment of the peptide predictive landscape through three key dimensions. First, we curated and standardized 15 high-quality datasets—totaling 161,571 unique sequences—by applying stringent biological filtering across diverse functional and physicochemical categories. Second, we conducted an extensive benchmarking of seven representative architectures, ranging from traditional machine learning with expert-crafted features to state-of-the-art pre-trained protein language models. Third, we performed a critical investigation into data partitioning strategies, specifically examining the efficacy of homology-based clustering via tools like MMseqs2. Notably, our analysis reveals intrinsic challenges in peptide clustering—such as high singleton ratios and uneven cluster distributions—which lead to a surprising convergence in model performance between random and similarity-based splits. To support community efforts, we provide the PPB online platform (http://ppb.molmatrix.com/index.html), offering centralized access to standardized datasets, interactive visualizations, and detailed evaluation protocols. Collectively, PPB not only offers a robust benchmarking resource but also highlights the unique methodological hurdles in evaluating peptide property prediction.

## Materials and methods

### Datasets in PPB

All datasets included in PPB are collected from publicly accessible peptide databases or previously published studies, covering representative application scenarios in peptide biological and physicochemical property prediction. Each dataset is fully traceable to its original source, ensuring data transparency and experimental reproducibility. In terms of task formulation, PPB comprises both classification and regression problems. To ensure comparability among different datasets and the reliability of evaluation results, all datasets were subjected to a unified data cleaning and standardization procedure before being incorporated into PPB. Specifically, samples containing non-standard amino acid characters or incomplete sequence information were removed, and fully duplicated sequences were merged. All datasets were then organized into a unified data format to ensure consistency in subsequent feature extraction, model training, and evaluation processes. This standardized processing effectively reduces potential biases introduced by differences in data quality. The detailed processing results of the classification and regression datasets are presented as follows.

#### AMP

The AMP (Antimicrobial peptide) dataset consists of positive samples collected from the APD3 [19], DBAASP [17], and DRAMP [20] databases. Only peptides with both N- and C-termini labeled as Free (or unlabeled) and composed exclusively of natural amino acids were retained, with a maximum sequence length of 183. Negative samples were obtained from the UniProt [21] database by filtering entries with subcellular location annotated as cytoplasm, restricting sequence length to fewer than 183 residues, and removing any entries containing keywords related to antimicrobial activity, including antimicrobial, antibiotic, antiviral, antifungal, effector, and excreted [22]. After unified preprocessing, the AMP dataset contains 47,511 sequences, including 28,750 positive samples and 18,761 negative samples, with sequence lengths ranging from 1 to 183 residues.

#### ASSEM

The ASSEM (Self-assembling peptide) dataset was collected from two published studies [23, 24]. After preprocessing, it contains 41,703 sequences, with 15,007 self-assembling peptides and 26,696 non-self-assembling peptides, and sequence lengths ranging from 3 to 24 residues.

#### CPP

The CPP (Cell-penetrating peptide) dataset was derived from the curated CPP924 dataset reported in [25]. After preprocessing, the dataset consists of 924 sequences, including 462 positive and 462 negative samples, with sequence lengths ranging from 4 to 61 residues.

#### HEMO, HUMAN, and SOLU

These datasets were collected from [26].After unified preprocessing, the HEMO (Hemolytic peptide) dataset contains 6,076 peptide sequences, including 1,311 hemolytic peptides and 4,765 non-hemolytic peptides, with sequence lengths ranging from 1 to 190 residues. The HUMAN (non-fouling peptide) dataset consists of 17,180 sequences, including 3,600 positive samples and 13,580 negative samples, with sequence lengths ranging from 5 to 198 residues. The SOLU (solubility peptide) dataset contains 18,453 peptide sequences, comprising 8,785 soluble peptides and 9,668 insoluble peptides, with sequence lengths ranging from 4 to 198 residues.

#### TOXIC

The TOXIC (Toxic peptide) dataset was obtained from the ToxinPred2 [27] database, retaining peptides with sequence lengths shorter than 200. After preprocessing, the dataset includes 1,516 sequences, with 1,052 toxic and 464 non-toxic peptides, and sequence lengths ranging from 35 to 200 residues.

#### AOPP

The AOPP (Antioxidant peptide) dataset was collected from the Antioxidant Peptide Prediction database [28]. After preprocessing, it contains 3,154 sequences, including 1,586 antioxidant peptides and 1,568 non-antioxidant peptides, with sequence lengths ranging from 2 to 20 residues.

#### NEU

The NEU (Neuropeptide) dataset was compiled from [29]. After preprocessing, it consists of 8,699 sequences, with balanced class distribution and sequence lengths ranging from 4 to 99 residues.

#### ADP

The ADP (Anti-diabetic peptide) dataset was collected from [30]. After preprocessing, it contains 5,667 sequences, with 418 positive and 5,249 negative samples, and sequence lengths ranging from 5 to 41 residues.

#### DPPIV

The DPPIV (Dipeptidyl peptidase-IV inhibitory peptide) dataset was obtained from [31]. After preprocessing, it includes 1,322 sequences, with balanced classes and sequence lengths ranging from 2 to 90 residues.

#### BBB

The BBB (Blood–brain barrier penetrating peptide) dataset was collected from [32], integrating the SCMB3PP training dataset 1 and test dataset 1. After preprocessing, it contains 521 sequences, including 265 BBB-penetrating peptides and 256 non-penetrating peptides, with sequence lengths ranging from 5 to 30 residues.

#### EC and SA

The EC (Escherichia coli inhibitory activity) and SA (Staphylococcus aureus inhibitory activity) datasets were collected from [33]. After preprocessing, the EC dataset contains 3,854 peptide sequences with a maximum sequence length of 60 residues, while the SA dataset consists of 3,152 sequences with a maximum sequence length of 60 residues.

#### HEMO reg

The HEMO (Hemolytic activity) regression dataset was obtained from the HemoPI2 database [34]. After preprocessing, the dataset comprises 1,839 peptide sequences with a maximum length of 39 residues, alongside their corresponding hemolytic concentrations (µM).

### Split methods in PPB

To systematically evaluate model performance under different generalization scenarios, PPB adopts two data splitting strategies: random splitting and sequence similarity–based splitting. For random splitting, samples are randomly divided into training, validation, and test sets with a fixed ratio of 8:1:1. For sequence similarity–based splitting, sequence clusters are used as the minimum splitting unit to explicitly control sequence similarity between subsets. Specifically, we employ MMseqs2 to cluster the entire sequence space and perform an extensive grid search over a predefined parameter landscape to optimize partitioning outcomes. This search space encompasses 36 distinct configurations, generated by varying the sequence identity threshold (–min-seq-id) and alignment coverage (-c) from 0.3 to 0.8 (with a step size of 0.1). High alignment sensitivity (-s 7) is consistently applied to address the intrinsic challenges of short peptide sequence alignment. We utilized the coverage mode 5 (–cov-mode 5), which calculates coverage based solely on the shorter sequence in each pair. This specific configuration is critical for peptide benchmarking as it prevents the artificial inflation of similarity scores caused by significant length disparities, ensuring that the alignment remains representative of the peptide’s core functional motifs. Following this systematic sweep, the resulting clusters are randomly shuffled and assigned to the training, validation, and test sets in an 8:1:1 ratio.

### Feature Representation in PPB

This study adopts multiple feature representation strategies according to the input requirements of different modeling paradigms, aiming to comprehensively evaluate the suitability of various representations for peptide property prediction tasks. In general, feature representations in PPB can be categorized into two types: handcrafted feature–based representations and end-to-end representations based on raw peptide sequences.

For traditional machine learning models, PPB employs handcrafted feature representations that map peptide sequences into fixed-dimensional numerical vectors. Specifically, these features [35] include amino acid composition frequencies, physicochemical descriptors, and one-hot encoding representations. Amino acid frequency features characterize the overall compositional properties of peptides by calculating the relative occurrence of each amino acid type. Physicochemical descriptors [36] capture global molecular properties of peptides, such as hydrophobicity, charge distribution, aliphatic index, and related statistical measures (see Table 1 in supplementary material). One-hot encoding explicitly represents amino acid identities while preserving positional information along the sequence. In PPB, all traditional machine learning models are trained and evaluated using the same feature sets and preprocessing procedures, ensuring that performance differences among algorithms primarily arise from the models themselves rather than biases introduced by feature selection.

For deep learning models and pretrained protein language models, PPB adopts sequence-based end-to-end representations. Unlike handcrafted features, these approaches take peptide sequences directly as input and automatically learn high-level representations from the raw sequences. In deep learning models, peptide sequences are first converted into integer-encoded representations, where each amino acid is mapped to a unique integer identifier. The encoded sequences are then projected into a continuous vector space through an embedding layer, followed by sequence modeling modules that learn contextual relationships among amino acids, thereby producing sequence representations for downstream prediction tasks. For pretrained protein language models, raw amino acid sequences are directly fed into the model. These models are pretrained on large-scale protein sequence corpora using self-supervised learning objectives, enabling them to capture rich semantic and structure-related information. In PPB, the output representations of pretrained models are used for peptide property classification or regression, allowing evaluation of the transferability of their general-purpose sequence representations to peptide-specific tasks.

### MODELS and Evaluation Protocol in PPB

To comprehensively evaluate the performance of different modeling paradigms in peptide property prediction tasks, PPB compares three representative categories of models: traditional machine learning models (ML), deep learning models (DL), and pretrained protein language models. Traditional machine learning models include Random Forest (RF), Support Vector Machine (SVM), and XGBoost (XGB), which take handcrafted sequence features as input. Deep learning models adopt sequence-based LSTM and Transformer encoders to automatically learn representations from peptide sequences through end-to-end training. Pretrained models include ESM [37, 38] and PEPBERT [39], which take raw amino acid sequences as input, keep the pretrained model parameters frozen, and perform classification or regression through task-specific multilayer perceptron (MLP) prediction heads. These models cover the major methodological paradigms commonly used in peptide property prediction, enabling systematic analysis of performance differences under a unified evaluation setting.

All models are compared under unified data splits, training procedures, and evaluation protocols to ensure fairness and reproducibility. For classification tasks, the model outputs binary values; for regression tasks, it directly predicts continuous values. Traditional machine learning models are implemented using scikit-learn, whereas deep learning models and pretrained models are implemented based on PyTorch. Model optimization is performed on the training set, with early stopping and model selection conducted on the validation set. All experiments are repeated under three fixed random seed settings to assess the stability of model performance and to mitigate the impact of randomness on the results. The specific hyperparameter configurations for all models are detailed in Supplementary Table 2 and Supplementary Table 3.

As for evaluation metrics, PPB adopts task-specific metrics to assess model performance across different task types. For classification tasks, Accuracy, Precision, Recall, and F1-score are used as evaluation metrics, with F1-score serving as the primary metric for model comparison to better reflect predictive performance under class-imbalanced settings. For regression tasks, Root Mean Squared Error (RMSE) and the coefficient of determination (R^2^) are employed to evaluate prediction error and goodness of fit, with RMSE used as the primary evaluation metric. Log-transformation was applied to continuous regression labels to mitigate heavy-tailed label distributions and to stabilize model training and evaluation. All evaluation metrics are computed on the independent test set, and the final reported results are obtained by averaging over multiple repeated experiments.

## Results

### Statistical Characterization of the PPB Datasets

The PPB framework encompasses 15 widely recognized public datasets, meticulously curated to span a diverse landscape of peptide functional activities. This collection includes 12 classification tasks—targeting antimicrobial (AMP), self-assembly (ASSEM), cell-penetrating (CPP), hemolytic (HEMO), antifouling (HUMAN), solubility (SOLU), toxicity (TOXIC), antioxidant (AOPP), neuroactive (NEU), antidiabetic (ADP), enzyme inhibitory (DPPIV) activities, and blood-brain barrier (BBB) permeability—and 3 regression tasks focused on quantifying anti-E. coli (EC), anti-S. aureus (SA), and hemolytic (HEMO reg) potencies. As summarized in Fig 2, these datasets exhibit substantial divergence in sequence scale, length distribution, isoelectric point (*pI*) profiles, and class proportions, providing a multifaceted foundation for evaluating model adaptability across distinct biological contexts.

**Fig 1.**
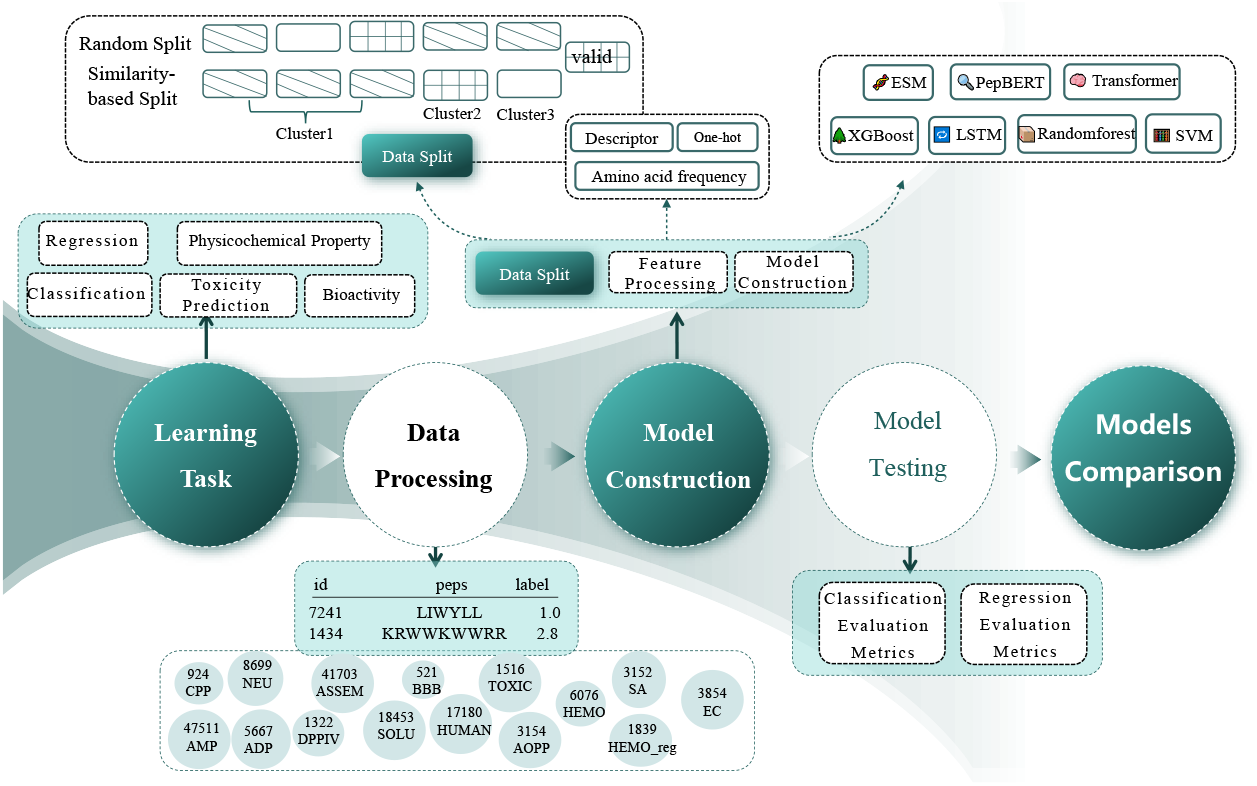
Schematic overview of the proposed comprehensive benchmarking framework for peptide property prediction. The pipeline comprises five sequential stages: Learning Tasks; Data Processing; Model Construction; and finally Model Testing and Models Comparison to evaluate performance via specific metrics, establishing standardized baselines for property prediction tasks.

**Fig 2.**
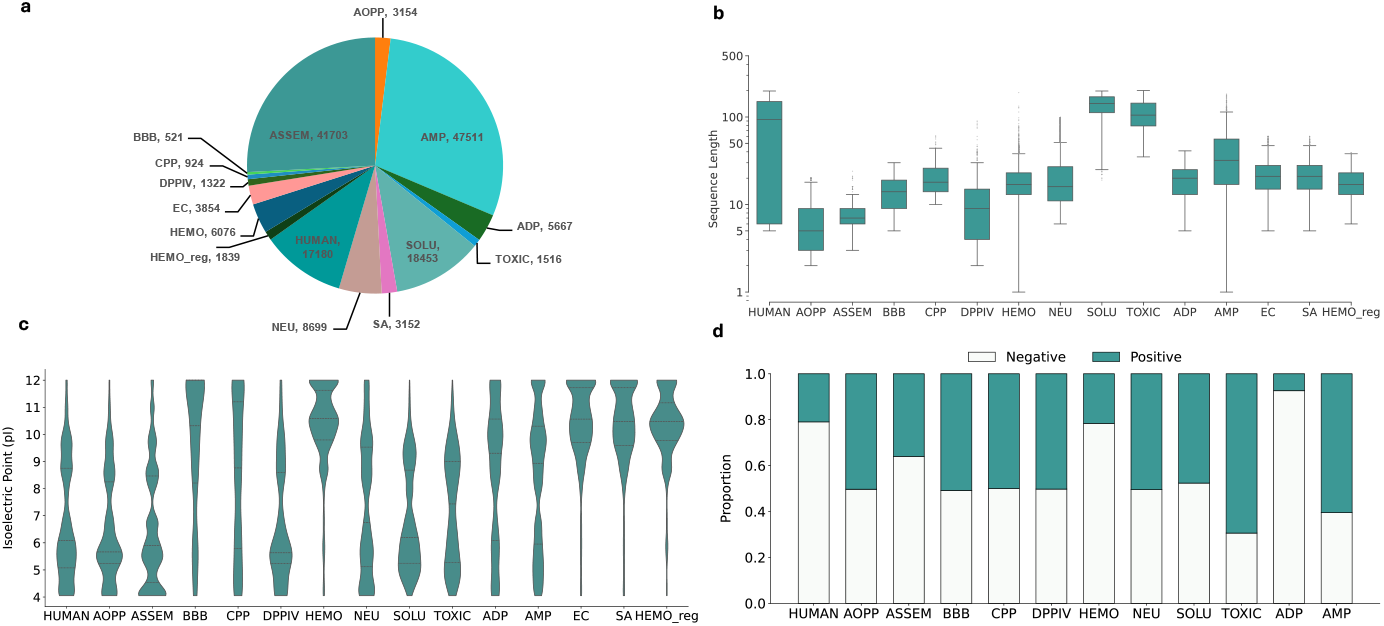
Statistical overview of the benchmark datasets used in the experiments. (a) Distribution of sequence counts across the benchmark datasets; (b) Distribution of peptide sequence lengths; (c) isoelectric point (pI) distributions; (d) class ratio distributions across classification datasets.

As illustrated in Fig 2 a–c, the benchmark is characterized by pronounced statistical heterogeneity. Regarding dataset scale, while the AMP, ASSEM, SOLU, and HUMAN datasets are large-scale (each exceeding 10,000 sequences), the remaining tasks are relatively low-resource, reflecting the data-scarcity challenges common in specific functional domains. Sequence length distributions reveal clear task-specific stratification: AOPP, ASSEM, and DPPIV are primarily dominated by short-chain oligopeptides (*<* 10 residues), whereas the majority of tasks (BBB, CPP, HEMO, NEU, ADP, AMP, EC, and SA) fall within the canonical peptide range of 10–50 residues. Notably, HUMAN, SOLU, and TOXIC datasets contain significantly longer sequences, with median lengths approaching or exceeding 100 residues and exhibiting broader variance. The *pI* distributions further highlight the diverse physicochemical signatures across functional classes; for instance, AMP and HEMO datasets show a distinct alkaline bias (*pI* peaks between 10–12), whereas most other datasets lean toward weakly acidic profiles, reflecting the specialized charge requirements for different biological functions.

Furthermore, the class ratio distributions (Fig 2 d) underscore the structural complexity of PPB. While tasks such as AOPP, BBB, CPP, and DPPIV maintain near-balanced distributions (1:1), others—most notably ADP, HEMO, and HUMAN—exhibit significant class imbalance, with the ADP dataset characterized by a starkly long-tailed distribution dominated by negative samples. This spectrum of class structures, from balanced to highly skewed, provides a rigorous testbed for assessing model robustness and the efficacy of various loss functions or sampling strategies in addressing real-world data irregularities.

### Impact of Feature Representation and Model Choice on Peptide Prediction Performance

To establish a robust baseline for peptide property prediction, we systematically evaluated 13 model configurations, comprising three traditional machine learning (ML) models (RF, SVM, and XGB) paired with three encoding schemes (one-hot, frequency, and physicochemical descriptors), two deep learning (DL) architectures (LSTM and Transformer), and two protein language models (pLMs: PepBERT and ESM). The aggregate performance across all classification and regression tasks is summarized in Fig 3 a and b. The detailed numerical performance metrics for all evaluated models are provided in Supplementary Table 4 and Supplementary Table 5.

**Fig 3.**
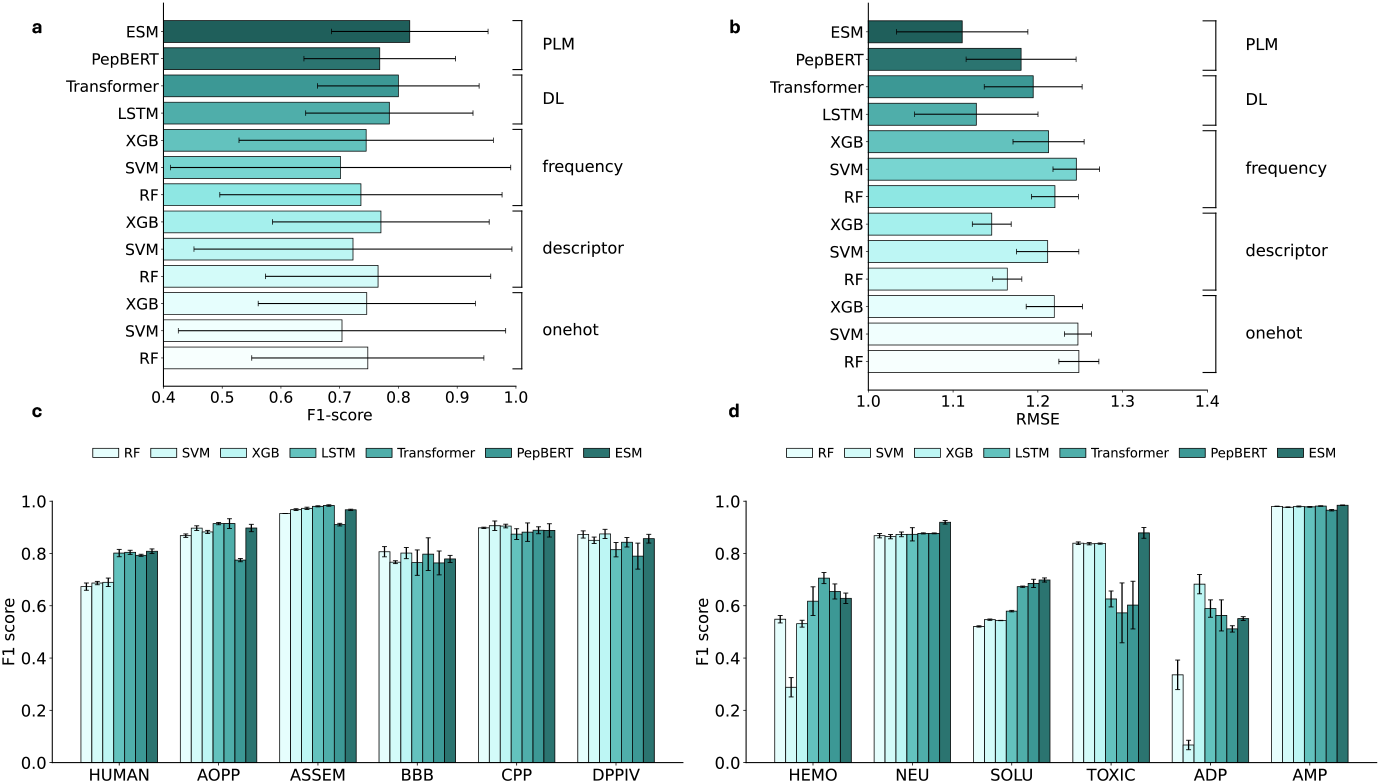
Average performance across datasets. (a) Average F1-score across 12 classification datasets for each model. (b) Average RMSE across 3 regression datasets for each model. (c-d) Bar charts showing the F1 scores of the machine learning model, two deep learning models, and two pretrained protein language models across 12 classification datasets. Bars represent the mean performance over repeated runs, with error bars indicating the standard deviation.

We first examined the impact of feature encoding on traditional ML models. Among the three tested representations, physicochemical descriptors consistently outperformed both one-hot encoding and amino acid composition frequency. This superiority underscores that descriptors, by integrating intrinsic biochemical properties, capture more functionally relevant information than schemes restricted to raw sequence composition. However, even with optimized descriptors, ML models generally lagged behind the DL architectures (LSTM and Transformer). This performance gap likely arises because, while descriptors aggregate global or local physicochemical properties, they often forfeit the precise spatial arrangement and sequential order of amino acids. In contrast, DL models inherently possess superior feature extraction capabilities, enabling them to learn complex, order-dependent patterns directly from the primary sequences.

Notably, the protein language models, particularly ESM, achieved the highest average F1 scores and the lowest RMSE values across the benchmark. Beyond absolute accuracy, ESM demonstrated significantly lower performance variance in classification tasks, indicating superior stability across diverse biological contexts. A granular analysis of specific datasets further highlights the advantage of pLMs; as shown in Fig 3 d and Fig 4, ESM exhibited markedly better performance on small-scale datasets such as NEU and SOLU, as well as regression tasks like SA, EC, and HEMO. This resilience to data scarcity likely stems from the knowledge transfer inherent in large-scale pre-training: by learning universal structural and functional grammars from billions of protein sequences, ESM can effectively regularize its predictions even when supervised downstream data are limited.

**Fig 4.**
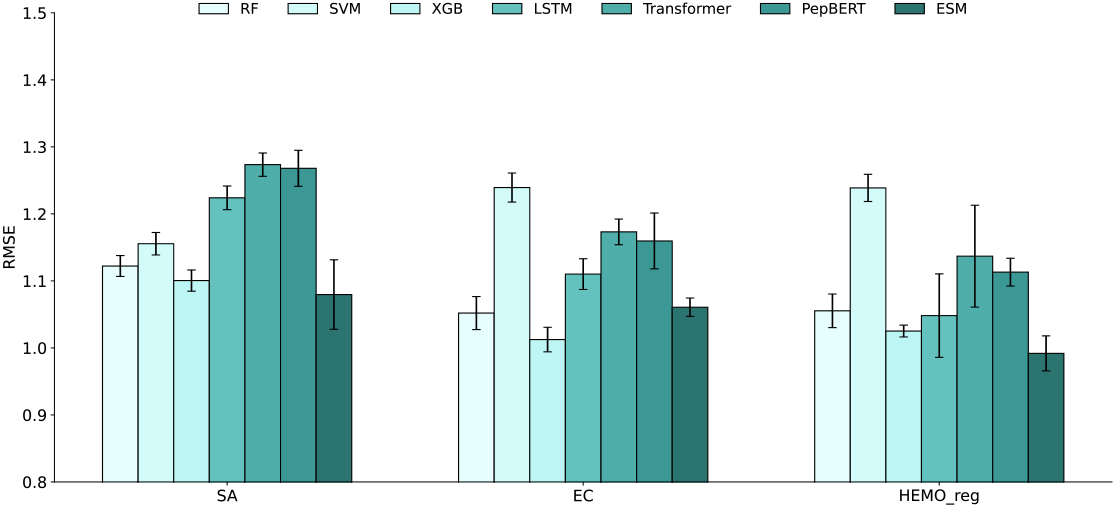
Bar charts showing the RMSE of the machine learning model, two deep learning models, and two pretrained protein language models across 3 regression datasets. Bars represent the mean performance over repeated runs, with error bars indicating the standard deviation.

From a data-centric perspective, we observed that model performance is heavily influenced by dataset composition. Tasks characterized by severe class imbalance, such as HEMO and ADP, yielded consistently lower performance metrics compared to more balanced datasets. This trend suggests that while advanced architectures like pLMs can mitigate small-sample issues, the intrinsic bias introduced by skewed label distributions remains a significant hurdle, necessitating more specialized sampling or loss-weighting strategies in future peptide design pipelines.

### Impact of Data Splitting Strategies on Model Generalization

To assess the efficacy of homology-based evaluation, we conducted a systematic exploration of similarity-based partitioning hyperparameters. For each dataset, we optimized combinations of sequence identity (*identity* ∈ [0.3, 0.8]) and alignment coverage (*coverage* ∈ [0.3, 0.8]) to identify configurations yielding the most robust clustering. However, our results reveal that even after dataset-specific tuning, similarity-based clustering remains highly fragmented and fails to establish the stable, representative groupings necessary for a rigorous evaluation of generalization.

As shown in Fig 5 a, the Gini coefficients for cluster sizes across all datasets are consistently below 0.5. While this initially suggests a lack of extreme imbalance, this low metric—in the context of peptide data—actually reflects a “pseudo-uniform” distribution caused by excessive fragmentation rather than meaningful structural organization. This is further corroborated by Fig 5 b, which shows a persistently high singleton ratio. Even in optimal cases where the cluster-to-sequence ratio approaches 0.5, the singleton ratio only marginally decreases to approximately 0.4. Moreover, Fig 5 c illustrates that the average cluster size remains below 3, with the 95th percentile staying within single digits. The detailed evaluation of various MMseqs2 clustering configurations and their impact on data partitioning is provided in Supplementary Table Collectively, these findings indicate that similarity-based clustering is dominated by a vast number of small clusters and isolated sequences. Such structures fail to form the cohesive “homology groups” required to meaningfully simulate unseen sequence space.

**Fig 5.**
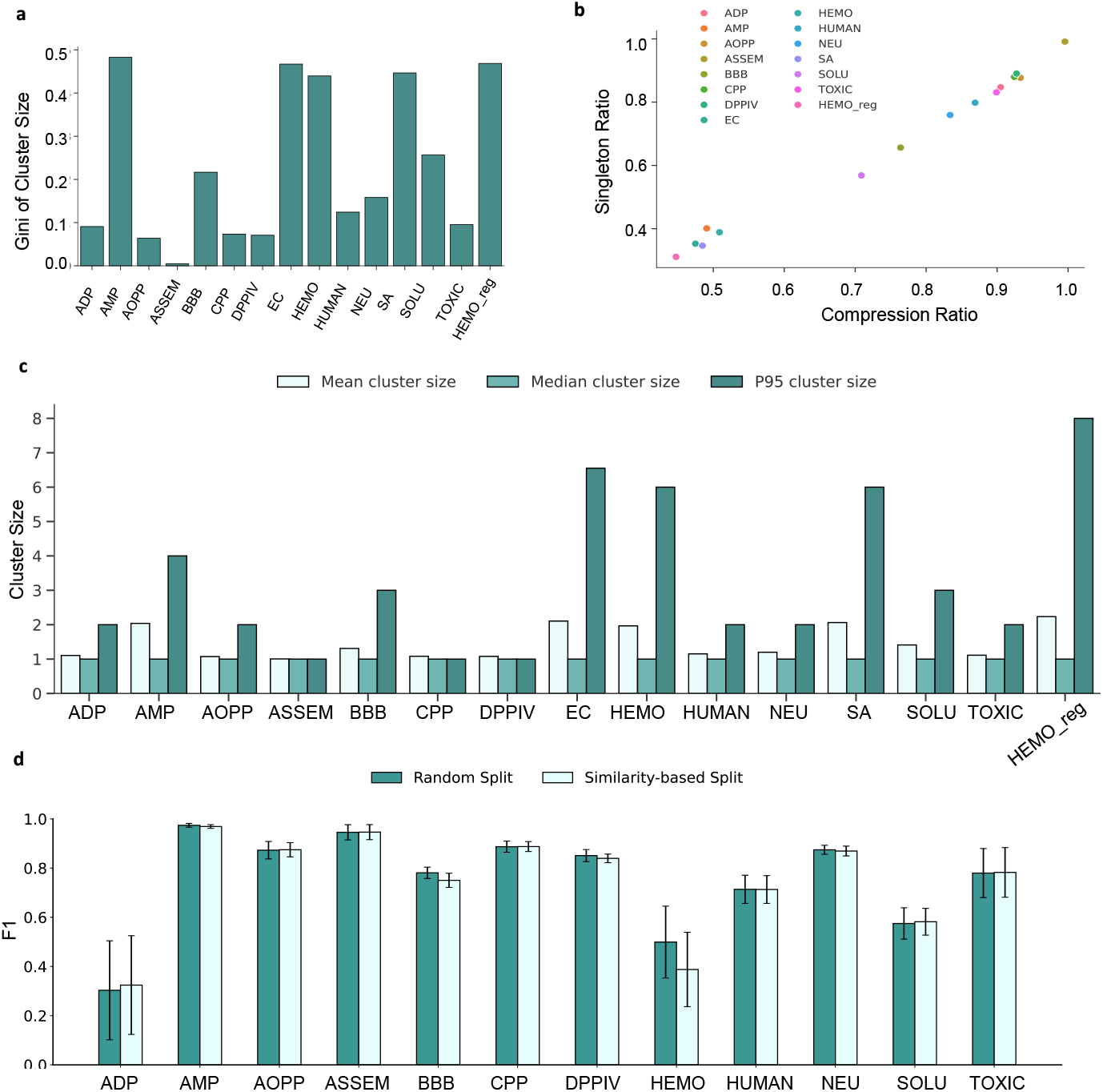
Evaluation of model robustness and generalization under different data splitting strategies. (a) Gini coefficient of cluster size distributions across datasets. (b) Relationship between clustering granularity and singleton ratio. (c) Cluster sizes after similarity-based clustering across different datasets.P95 cluster size represents the size threshold that covers 95% of the cluster’s data units. (d) Average F1 scores on 12 classification datasets under different data splitting strategies.

**Fig 6.**
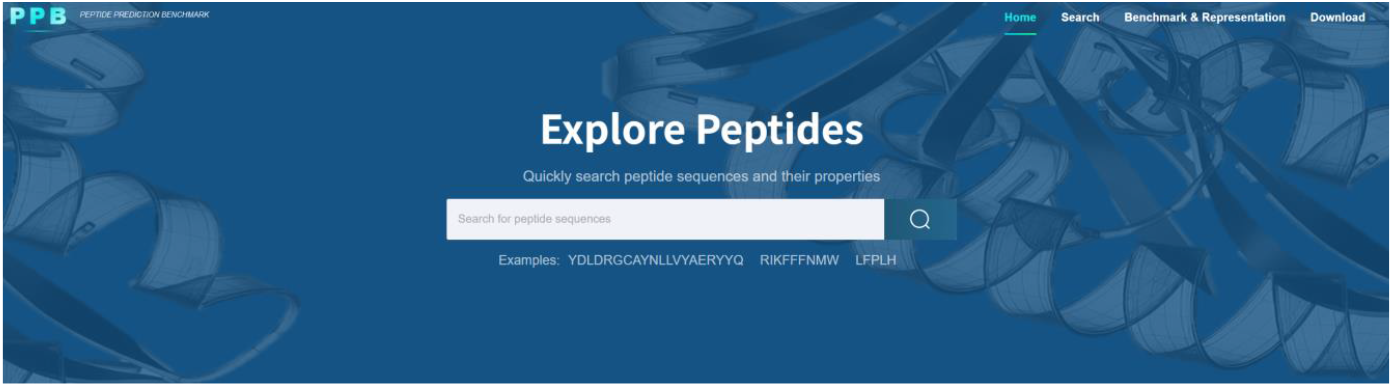
Overview of the benchmark website developed for peptide property prediction. The figure shows the homepage interface of the online platform, which provides unified access to benchmark datasets, data splitting strategies, and model evaluation results presented in this study.

Building on this structural analysis, we compared model performance under random versus similarity-based splitting strategies. As illustrated in Fig 5 d, the performance metrics under both regimes are remarkably consistent. Contrary to expectations in protein-level modeling, similarity-based splitting did not yield more conservative or discriminative performance estimates. We attribute this phenomenon to the inherent limitations of current clustering algorithms (e.g., MMseqs2) when applied to short peptide sequences; because these tools fail to produce well-structured partitions, the resulting train-test splits do not significantly differ from random partitions in terms of their effective data distribution. Consequently, our benchmark suggests that for the current scale and sequence characteristics of peptide datasets, random splitting provides a practical and sufficient estimate of performance, while the benefits of traditional homology-based partitioning remain constrained by the difficulty of defining meaningful clusters in short-sequence space.

### PPB Web Server

To improve the accessibility and reproducibility of the PPB benchmark, we developed an online platform named PPB (available at: http://ppb.molmatrix.com/index.html) for centralized dissemination of the benchmark resources. The website provides unified access to the unsplit datasets, randomly split datasets, and sequence similarity–based split datasets included in PPB, enabling consistent evaluation under different data partitioning strategies. In addition, the platform offers visualizations of benchmark results for both classification and regression tasks, together with a brief introduction to the benchmark construction and evaluation protocols. By integrating data access and result presentation into a single interface, the PPB web server facilitates systematic model comparison and practical usage of the PPB benchmark.

## Discussion

While this benchmark provides a systematic evaluation of various architectures and feature representations, several critical challenges remain that point toward future research directions. A primary limitation of the current study is the reliance on primary sequence information, without explicitly incorporating three-dimensional (3D) structural features or conformational dynamics. Unlike folded proteins with relatively stable scaffolds, peptides are characterized by high conformational flexibility and often undergo induced fit transitions upon binding to their biological targets. Their functional properties—such as cell permeability and binding affinity—are governed not only by the amino acid sequence but also by the spatial distribution of pharmacophores and the stability of specific secondary structures (e.g., *α*-helices or *β*-turns). Consequently, sequence-only representations may reach a “performance ceiling” in tasks where spatial configuration and molecular interactions are the dominant determinants of activity.

To overcome these limitations, future modeling efforts could pivot toward structure-aware or physics-informed learning. With the advancement of high-accuracy structure prediction tools like AlphaFold3 and Boltz, integrating predicted ensembles of peptide conformations into the learning pipeline has become increasingly feasible. Furthermore, incorporating physical constraints (e.g., electrostatic maps or solvent accessibility) and experimental priors (e.g., NMR-derived constraints) could bridge the gap between static sequence modeling and dynamic functional reality.

Beyond structural considerations, our findings regarding the inefficacy of current clustering algorithms on short peptides underscore a methodology gap in the field. The high fragmentation observed in similarity-based splitting suggests that “homology” in the peptide world may need to be redefined—perhaps shifting from global sequence alignment to motif-based or pharmacophore-based similarity. Developing such specialized partitioning strategies will be essential for a more rigorous assessment of model extrapolation in unseen chemical spaces.

Ultimately, the PPB benchmark serves as a foundational step. By highlighting the strengths of protein language models and the unique hurdles in peptide data organization, it sets the stage for the next generation of predictive tools that are not only sequence-savvy but also structurally and physically grounded.

## Acknowledgments

This work was supported by the National Natural Science Foundation of China (Grants 62572374, U22A2037, and 62132015).

## References

1. Markus Muttenthaler DJA Glenn F King, Alewood PF. Trends in peptide drug discovery. Nature Reviews Drug Discovery. 2021;20:309–25.

2. Wang L, Wang N, Zhang W, Cheng X, Yan Z, Shao G, et al. Therapeutic peptides: current applications and future directions. Signal Transduction and Targeted Therapy. 2022;7(1):48.

3. Mallick P, Schirle M, Chen SS, Flory MR, Lee H, Martin D, et al. Computational Prediction of Proteotypic Peptides for Quantitative Proteomics. Nature Biotechnology;25(1):125–31. Available from: https://doi.org/10.1038/nbt1275. doi:10.1038/nbt1275.

4. Lin C, Xiong S, Cui F, Zhang Z, Shi H, Wei L. Deep Learning in Antimicrobial Peptide Prediction. Journal of Chemical Information and Modeling;65(14):7373–92. Available from: https://doi.org/10.1021/acs.jcim.5c00530. doi:10.1021/acs.jcim.5c00530.

5. Chen X, Li C, Bernards MT, Shi Y, Shao Q, He Y. Sequence-based peptide identification, generation, and property prediction with deep learning: a review. Molecular Systems Design & Engineering. 2021;6(6):406–28.

6. Lei Y, Li S, Liu Z, Wan F, Tian T, Li S, et al. A deep-learning framework for multi-level peptide–protein interaction prediction. Nature communications. 2021;12(1):5465.

7. Yan J, Bhadra P, Li A, Sethiya P, Qin L, Tai HK, et al. Deep-AmPEP30: improve short antimicrobial peptides prediction with deep learning. Molecular Therapy Nucleic Acids. 2020;20:882–94.

8. Olsen TH, Yesiltas B, Marin FI, Pertseva M, García-Moreno PJ, Gregersen S, et al. AnOxPePred: using deep learning for the prediction of antioxidative properties of peptides. Scientific reports. 2020;10(1):21471.

9. Lin C, Xiong S, Cui F, Zhang Z, Shi H, Wei L. Deep learning in antimicrobial peptide prediction. Journal of Chemical Information and Modeling. 2025;65(14):7373–92.

10. Veltri D, Kamath U, Shehu A. Deep learning improves antimicrobial peptide recognition. Bioinformatics. 2018;34(16):2740–7.

11. Zeng WF, Zhou XX, Willems S, Ammar C, Wahle M, Bludau I, et al. AlphaPeptDeep: a modular deep learning framework to predict peptide properties for proteomics. Nature Communications. 2022;13(1):7238.

12. Zhang H, Saravanan KM, Wei Y, Jiao Y, Yang Y, Pan Y, et al. Deep learning-based bioactive therapeutic peptide generation and screening. Journal of Chemical Information and Modeling. 2023;63(3):835–45.

13. Chen J, Cheong HH, Siu SW. xDeep-AcPEP: deep learning method for anticancer peptide activity prediction based on convolutional neural network and multitask learning. Journal of chemical information and modeling. 2021;61(8):3789–803.

14. Al-Omari AM, Akkam YH, Zyout A, Younis S, Tawalbeh SM, Al-Sawalmeh K, et al. Accelerating antimicrobial peptide design: Leveraging deep learning for rapid discovery. Plos one. 2024;19(12):e0315477.

15. Chou KC. Prediction of protein cellular attributes using pseudo-amino acid composition. PROTEINS. 2001;43(3):246–55.

16. Daniel Veltri AS Uday Kamath. Deep learning improves antimicrobial peptide recognition. Bioinformatics. 2018;34(16):2740–7.

17. Pirtskhalava M, Amstrong AA, Grigolava M, Chubinidze M, Alimbarashvili E, Vishnepolsky B, et al. DBAASP v3: database of antimicrobial/cytotoxic activity and structure of peptides as a resource for development of new therapeutics. Nucleic Acids Research. 2021;49(D1):D288–97.

18. Rao R, Bhattacharya N, Thomas N, Duan Y, Chen P, Canny J, et al. Evaluating Protein Transfer Learning with TAPE. In: Advances in Neural Information Processing Systems. vol. 32. Curran Associates, Inc.; 2019..

19. Guangshun Wang ZW Xia Li. APD3: the antimicrobial peptide database as a tool for research and education. Nucleic Acids Research. 2016;44(D1):D1087–93.

20. Shi G, Kang X, Dong F, Liu Y, Zhu N, Hu Y, et al. DRAMP 3.0: an enhanced comprehensive data repository of antimicrobial peptides. Nucleic acids research. 2022;50(D1):D488–96.

21. The UniProt Consortium. UniProt: the Universal Protein Knowledgebase in 2023. Nucleic Acids Research. 2023;51(D1):D523–31.

22. Ma Y, Guo Z, Xia B, Zhang Y, Liu X, Yu Y, et al. Identification of antimicrobial peptides from the human gut microbiome using deep learning. Nature Biotechnology. 2022;40(6):921–31.

23. Liu Z, Wang J, Luo Y, Zhao S, Li W, Li SZ. Efficient prediction of peptide self-assembly through sequential and graphical encoding. Briefings in Bioinformatics. 2023;24(6):bbad409.

24. Njirjak M, Žužić L, Babić M, Janković P, Otović E, Kalafatovic D, et al. Reshaping the discovery of self-assembling peptides with generative AI guided by hybrid deep learning. Nature machine intelligence. 2024;6(12):1487–500.

25. Xiangzheng Fu XZ Lijun Cai, Zou Q. StackCPPred: a stacking and pairwise energy content-based prediction of cell-penetrating peptides and their uptake efficiency. Bioinformatics. 2020;36(10):3028–34.

26. Guntuboina C, Das A, Mollaei P, Kim S, Barati Farimani A. Peptidebert: A language model based on transformers for peptide property prediction. The Journal of Physical Chemistry Letters. 2023;14(46):10427–34.

27. Neelam Sharma SJ Leimarembi Devi Naorem, Raghava GPS. ToxinPred2: an improved method for predicting toxicity of proteins. Briefings in Bioinformatics. 2022;23(5):1–20.

28. Antioxidant Peptide Prediction (AOPP);. Accessed January 2025. http://inova.aibiochem.net/antioxpep/.

29. Wang L, Huang C, Wang M, Xue Z, Wang Y. NeuroPred-PLM: an interpretable and robust model for neuropeptide prediction by protein language model. Briefings in Bioinformatics. 2023;24(2):bbad077.

30. Yue J, Xu J, Li T, Li Y, Chen Z, Liang S, et al. Discovery of potential antidiabetic peptides using deep learning. Computers in Biology and Medicine. 2024;180:109013.

31. Charoenkwan P, Nantasenamat C, Hasan MM, Moni MA, Lio’ P, Manavalan B, et al. StackDPPIV: A novel computational approach for accurate prediction of dipeptidyl peptidase IV (DPP-IV) inhibitory peptides. Methods. 2022;204:189–98.

32. Phasit Charoenkwan CN, Hasan MM. Improved prediction and characterization of blood-brain barrier penetrating peptides using estimated propensity scores of dipeptides. Journal of Computer-Aided Molecular Design. 2022;36:781–96.

33. Cai J, Yan J, Un C, Wang Y, Campbell-Valois FX, Siu SWI. BERT-AmPEP60: A BERT-Based Transfer Learning Approach to Predict the Minimum Inhibitory Concentrations of Antimicrobial Peptides for Escherichia coli and Staphylococcus aureus. Journal of Chemical Information and Modeling. 2025;65(7):3186–202.

34. Chaudhary K, Kumar R, Singh S, Tuknait A, Gautam A, Mathur D, et al. A web server and mobile app for computing hemolytic potency of peptides. Scientific reports. 2016;6(1):22843.

35. Bonidia RP, Sampaio LDH, Domingues DS, Paschoal AR, Lopes FM, de Carvalho ACPLF, et al. Feature extraction approaches for biological sequences: a comparative study of mathematical features. Briefings in Bioinformatics. 2021;22(5):bbab011.

36. Özçelik R, van Weesep L, de Ruiter S, Grisoni F. peptidy: a light-weight Python library for peptide representation in machine learning. Bioinformatics Advances. 2025;5(1):vbaf058.

37. Rives A, Meier J, Sercu T, Goyal S, Lin Z, Liu J, et al. Biological structure and function emerge from scaling unsupervised learning to 250 million protein sequences. Proceedings of the National Academy of Sciences. 2021;118(15):e2016239118.

38. Lin Z, Akin H, Rao R, Hie B, Zhu Z, Lu W, et al. Evolutionary-scale prediction of atomic-level protein structure with a language model. Science. 2023;379(6637):1123–30.

39. Elnaggar A, Heinzinger M, Dallago C, Rehawi G, Wang Y, Jones L, et al. Prottrans: Toward understanding the language of life through self-supervised learning. IEEE transactions on pattern analysis and machine intelligence. 2021;44(10):7112–27.

